# Targeted long-read RNA sequencing reveals the complexity of *CLN3* transcription and the consequences of the most common 1-kb deletion in patients with juvenile CLN3 disease

**DOI:** 10.1101/2025.04.24.650398

**Authors:** Christopher J Minnis, Hao-Yu Zhang, Emil Gustavsson, Jasmaine Lee, Claire Anderson, Emily Gardner, Angela Schulz, Miriam Nickel, Gavin Arno, Neringa Jurkute, Andrew R Webster, Nicola Gammaldi, Filippo M Santorelli, Paul Gissen, Phillipa Mills, Mina Ryten, Sara E Mole

**Author notes:** Corresponding author: Christopher J. Minnis. Department of Biological and Environmental Sciences and Technologies (DiSTeBA), University of Salento, 73100 Lecce, Italy (Nicola Gammaldi).

## Abstract

Most genes are not yet fully annotated, and the extent of their transcript diversity and the roles and significance of specific isoforms is not understood. This information is therefore lacking for disease genes. The *CLN3* gene underlies classic juvenile CLN3 disease, also known as juvenile neuronal ceroid lipofuscinosis, a rare paediatric neurodegenerative disorder. The most common cause of this biallelic disorder is a 1-kb intragenic deletion that removes two internal coding exons (exons 7 and 8). Here, we report findings from the first long-read RNA sequencing targeting *CLN3* in blood samples derived from control individuals and from patients clinically and genetically diagnosed with juvenile CLN3 disease. We find that *CLN3* transcription is complex, with >80 different transcripts encoding >35 different open reading frames (ORF) of different lengths, and no dominantly expressed transcript. The 1-kb deletion has direct consequences on this. This is consistent across patients, with total loss of some transcripts including those encoding the canonical 438 amino acid protein and other significant smaller isoforms. The highest expressed disease transcripts include those lacking exons 7 and 8 and encoding a 181 amino acid protein isoform, and other novel isoforms that lack additional exons and encode longer ORFs. The different effects on transcription of other CLN3 disease-causing variants are revealed in single patients. Together, these findings confirm the complexity of transcription at the *CLN3* locus, reveal the impact of the 1-kb deletion and other variants on isoform abundance, and highlight the importance of understanding the contribution of these isoforms to CLN3 function in health and disease. Moreover, they impact the future design and development of personalised therapeutics and the design and generation of disease models. Finally, they underline the importance of full annotation for disease genes.

## Introduction

Transcript diversity has an important role in eukaryotes, with alternative splicing increasing the transcripts and proteins derived from a single coding gene. 94% of human genes undergo alternative splicing [1, 2] producing on average ∼4 transcripts per gene [3]. However, some genes produce numerous transcripts. This appears to be particularly true for genes expressed in the brain, which shows increased diversity of alternative spliced isoforms compared to other tissue types [2, 4]. The recognition and implication of alternative splicing in a disease context is becoming increasingly important for therapeutic development [5-7]. The effect of specific variants is particularly challenging to investigate for rare inherited brain diseases, unless patients share common disease alleles.

One of the most common types of Batten disease, a group of biallelic paediatric neurodegenerative disorders also known as the neuronal ceroid lipofuscinoses (NCL), is juvenile CLN3 disease resulting from variation in the gene *CLN3* [8]. The classic form of the disease accounts for ∼50% of all NCL cases, with first symptoms presenting at around 6-8 years old as acute visual loss (retinitis pigmentosa), followed by a progressive decline in cognitive and motor abilities, seizures and eventually premature death in early adulthood [8-10]. The canonical *CLN3* gene of 15 exons encodes a protein of 438 amino acids (here termed CLN3-438aa) [8]. The predominant mutation on one or both disease alleles in ∼70% of juvenile CLN3 patients is a 1-kb intragenic deletion that removes 966 bp between intron 6 to intron 8 (rs1555468634, g.28485965_28486930del, c.461-280_677+382del), causing the loss of two internal coding exons (7 and 8) of the CLN3 canonical transcript [11, 12].

Previous research on the consequences of this 1-kb deletion revealed the presence of two novel transcripts, at the time termed ‘major’ and ‘minor’ [13]; the ‘major’ transcript includes coding sequences from exons 1-6 before being spliced out of frame into coding exon 9, leading to a truncated protein of 181 amino acids due to a premature stop codon after 28 novel amino acids (here, CLN3-181aa); the ‘minor’ transcript contains a splice event from exon 6 to exon 10 that brings the coding sequence back into frame, leading to a longer protein of 328 amino acids containing the canonical N- and C-terminal residues and lacking 110 internal residues (here, CLN3-328aa). The NCL field had originally thought that the 1-kb deletion leads to a complete loss of function of CLN3, however, expressing the equivalents of these two transcripts in the fission yeast *Schizosaccharomyces pombe* suggested both retain some function [14] and a strain expressing the equivalent of the minor transcript was shown to retain functionality as well as gain novel functions unique to that sequence [15].

Recently, we examined the range of *CLN3* transcripts in publicly available untargeted long-read RNA-sequencing (lrRNAseq) databases [16]. This work, which analysed untargeted long-read data from ENCODE, highlighted that CLN3 annotation is incomplete and that there is a high variation of transcripts leading to many different isoformic open reading frames (ORF) that could theoretically be translated into different sized proteins [16]. In addition, we observed the ‘major’ transcript in healthy tissue samples (not affected by CLN3 disease), and drawing on public mass spectrometry data, provided evidence that this is translated into protein [16]. To address this knowledge gap in the study of *CLN3* transcription and to investigate the effect of variations in *CLN3* that cause disease, we set out to generate lrRNAseq data on *CLN3* transcripts from control individuals carrying no sequence variation in *CLN3* and from CLN3 disease patients homozygous or heterozygous for the disease-causing 1-kb intragenic deletion. This learning will be critical to the full understanding of CLN3 disease aetiology.

We now report the first study to investigate the full consequences of the 1-kb deletion in juvenile CLN3 disease patients at a transcriptional level. We used PacBio lrRNAseq technology that provides sequence along the full length of individual transcripts, and targeted the *CLN3* locus with hybridization probes to give a greater sequencing depth than a non-targeted approach. We identified the transcript profile of *CLN3* in 30 blood samples collected from CLN3 disease patients and control samples. This approach provides unparalleled insight into the transcript diversity of *CLN3* in healthy and disease states and allows us to speculate on the contribution of specific transcripts isoforms to disease and their therapeutic relevance.

## Results

### Targeted long-read RNA sequencing reveals many novel *CLN3* transcripts

To generate RNA for analysis of the consequences of genetic variation we use blood which can be easily and non-invasively sampled then transported at ambient temperatures before processing, enabling purification of RNA with high RNA integrity numbers (RIN). This is suitable for transcriptional analysis of *CLN3* because blood lymphocytes are affected by disease, evidenced by lysosomal storage material and the longstanding use of lymphocyte vacuolization as a marker phenotype of CLN3 disease patients [17], and our previous analysis of untargeted *CLN3* transcription across multiple tissues showing the largest diversity of transcripts in blood [16].

To provide the deepest mRNA sequence coverage we used specific hybridization probes that target *CLN3*, in contrast to extracting the sequences out of whole genome lrRNAseq data. We first confirmed the selectivity of this targeted technological approach using commercially available RNA (from Takara Bio) purified from six different tissues sourced from multiple healthy individuals (**Suppl Table 1**) and performing Pacific Biosciences (PacBio) lrRNAseq. Combining results from all samples (**Suppl Fig 1)**, a total of 590 different *CLN3* transcripts are identified across these tissues. All tissues tested, including blood, show similar numbers of *CLN3* transcripts, with most being classed as coding known or coding novel. There is some variation in the proportion of each class of transcript across tissues (**Suppl Fig 1C**), likely due to the tissues being a mix of many cell types. The data for 25 transcripts ranked by proportional usage are summarised in **Suppl Fig 1A’**. Nine different transcripts are present in all tissue samples (**Suppl Fig 1A”**), and there is no dominant *CLN3* transcript in any tissue. Two transcripts show expression >10%. Considering the proteins that are encoded by these transcripts, the data for the top 25 open reading frames (ORF) are shown **in Suppl Fig 1B’**. Five ORFs are encoded in all tissues (**Suppl Fig 1B’’)**. Three ORFs show expression >10%. These data are consistent with our previous description of the range of transcripts identified through untargeted lrRNAseq [16]. Together, these results confirm blood is a suitable tissue for targeted lrRNAseq of *CLN3*.

Confident in the selectivity, reliability and depth of data when targeting transcripts, and that blood provides excellent representation of *CLN3* full length transcript isoforms, we similarly performed lrRNAseq targeting the *CLN3* gene for 30 blood samples (**Suppl Table 2**). The individuals with CLN3 disease span a range of ages and include children and teenagers, which made it challenging to provide age-matched controls as such samples are generally only available from other patients. For control samples we therefore used adults and children affected by other genetically defined types of Batten disease, unaffected family members including children, and younger and older adults affected by other genetically defined retinal diseases as CLN3 disease presents with retinitis pigmentosa.

We found that *CLN3* is targeted with high confidence in blood tissue, and that the targeted transcript profiles for control blood samples produce the expected structures with many similarities to the targeted transcripts of the commercial tissues (**Suppl Fig 1**). The total number of *CLN3* structural transcripts identified in blood is 679, with 336 of these in control samples (no CLN3 disease) and 467 of these in CLN3 disease patients (note, some transcripts appear in both subgroups) (**Graphical figure**). It is notable that *CLN3* transcript complexity in patient samples is greater than in control samples, with a higher number of unique transcripts expressed at a low level (<5% proportionally) in CLN3 disease patients than in controls (**Suppl Fig 2**).

There are differences between the categories of *CLN3* transcripts for control and patient samples (**Fig 1**). Both data sets had a high proportion of transcripts predicted to be coding novel, as expected for a gene that is under-represented in annotation. For all control samples together most transcripts are classed as coding (43.75%), either coding known (4.17%) or coding novel (39.58%); few are predicted as NMD (31.25%), either NMD known (2.38%) or NMD novel (28.87%) and few are predicted as non-coding (25%) (**Graphical figure**), as for the transcripts described for commercial RNA (**Suppl Fig 1**). For all patients’ samples together, fewer transcripts are classed as coding (35.33%), either coding known (3.00%) or coding novel (32.33%); many more are predicted as NMD (47.97%), either NMD known (1.07%) or NMD novel (46.90%) and non-coding (16.7%) (**Graphical figure**). 313 novel transcripts were identified for wild-type *CLN3* out of the 336 detected, meaning 93.15% of transcripts are underrepresented within the public database. Within CLN3 disease patients 447 (95.72%) out of the 467 detected *CLN3* transcripts are classed as novel.

**Figure 1:**
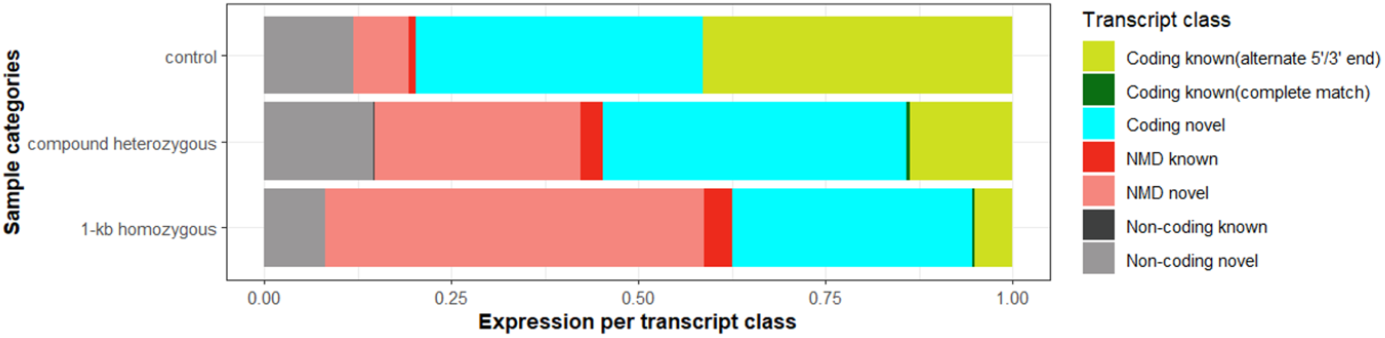
Overview summary of *CLN3* transcriptional data. Proportional expression of each transcript classification (coding known or novel, NMD known or novel, non-coding known or novel) per subgroup.

These data highlight the lack of knowledge around *CLN3* transcription in healthy individuals, as well as in CLN3 disease patients, and promise unprecedented detail in annotation, and especially the consequences of the 1-kb intragenic deletion. This is discussed in the next section.

### 47 novel protein-encoding *CLN3* transcripts are identified

We examined *CLN3* transcription in the 16 control individuals in more detail. The 336 transcripts consist of 84 unique structures (**Graphical figure**). Of these, only 11 (13%) transcripts are classed as coding known (3’/5’ alternative ends but known coding). 47 (56%) are classed as coding novel and are detected in all 16 control samples. These 47 transcripts encode 37 unique ORFs. Of the 26 detected transcripts predicted to undergo NMD, 5 (6%) are classed as known NMD transcripts and 21 (25%) as novel NMD transcripts.

Data on *CLN3* transcription is displayed for the top 25 ranked transcripts in **Fig 2A**, and for the top 10 encoded ORFs in **Fig 3A**. These groupings account for 75.1% of all detected transcripts and 95.6% of all ORF usage respectively. The top two *CLN3* transcripts, observed in all individuals, represent the coding known canonical ORF (CLN3-438aa) with significantly different alternative 5’UTRs. The highest expressed transcript (CLN3_6_438aa_5UTR_563_3UTR_386) has a median usage of ∼28% (range: 46.8 - 10.0)), and the next (CLN3_6_438aa_5UTR_290_3UTR_387) a median usage of ∼5.5% (range: 10.5 - 3.1). This confirms that there is no single dominant (i.e. >50% proportionally) transcript within blood tissue for *CLN3* expression (**Fig 2A**). Both of these transcripts are present in all control (**Suppl Fig 3**) and commercial tissue RNA samples (**Suppl Fig 1**). The canonical CLN3-438aa represents the highest expressed ORF in control samples, accounting for an average of 43.4% (range: 58.7-17.8) of all ORF (**Fig 3A**).

**Figure 2:**
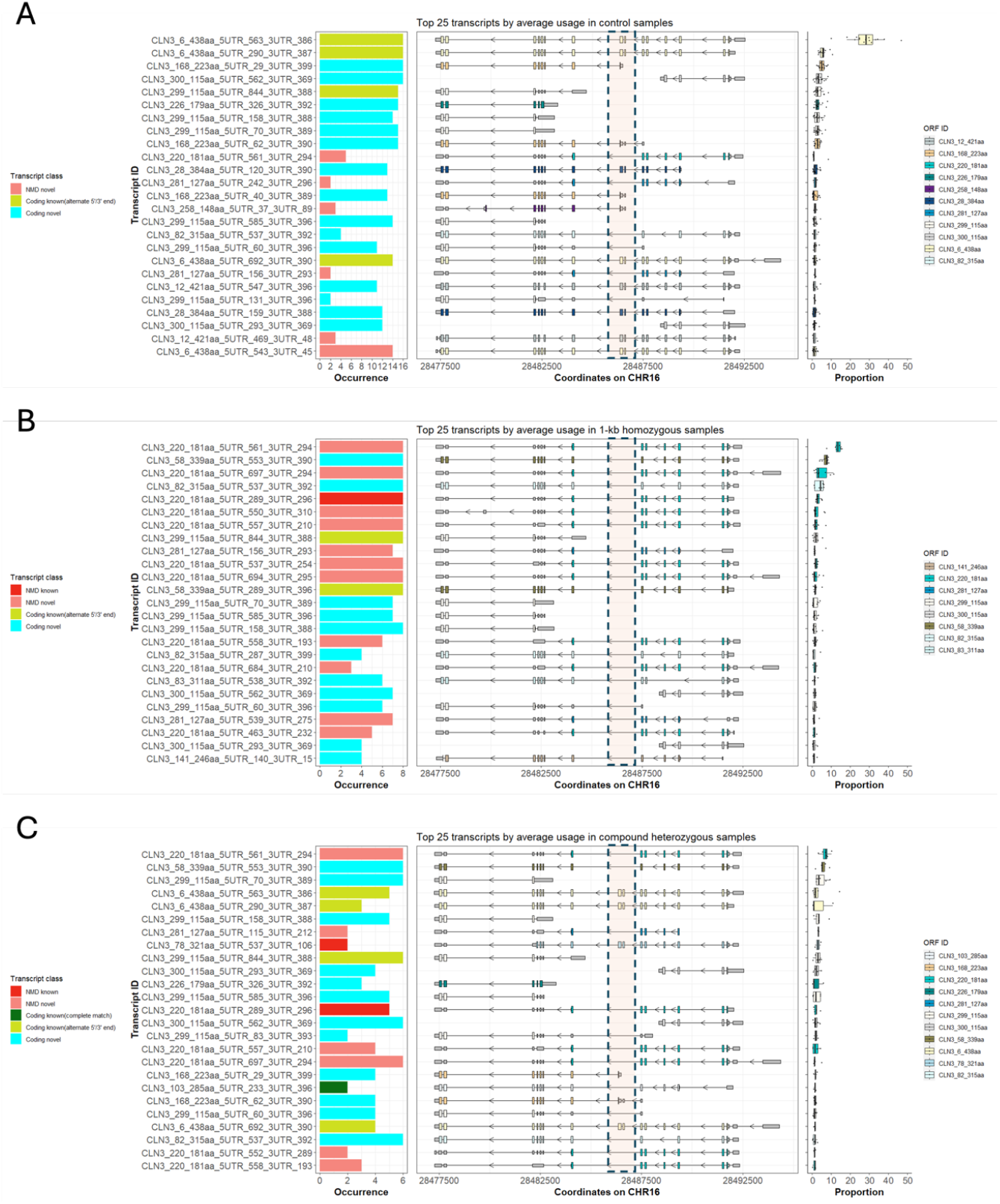
Overview of *CLN3* transcript diversity in controls and 1-kb deletion patients. A-C: From left to right, graphical representations of the top 25 transcripts in controls (A), homozygous patients (B), and compound heterozygous patients (C) by transcript occurrence (left panel), transcript structure with predicted open reading frames (ORF) (coloured extended boxed) and non-coding exons (grey, smaller boxes) (middle panel) and average transcript expression based on normalised usage (right panel). Approximate coordinates of the 1-kb deletion are indicated by salmon boxes.

**Figure 3:**
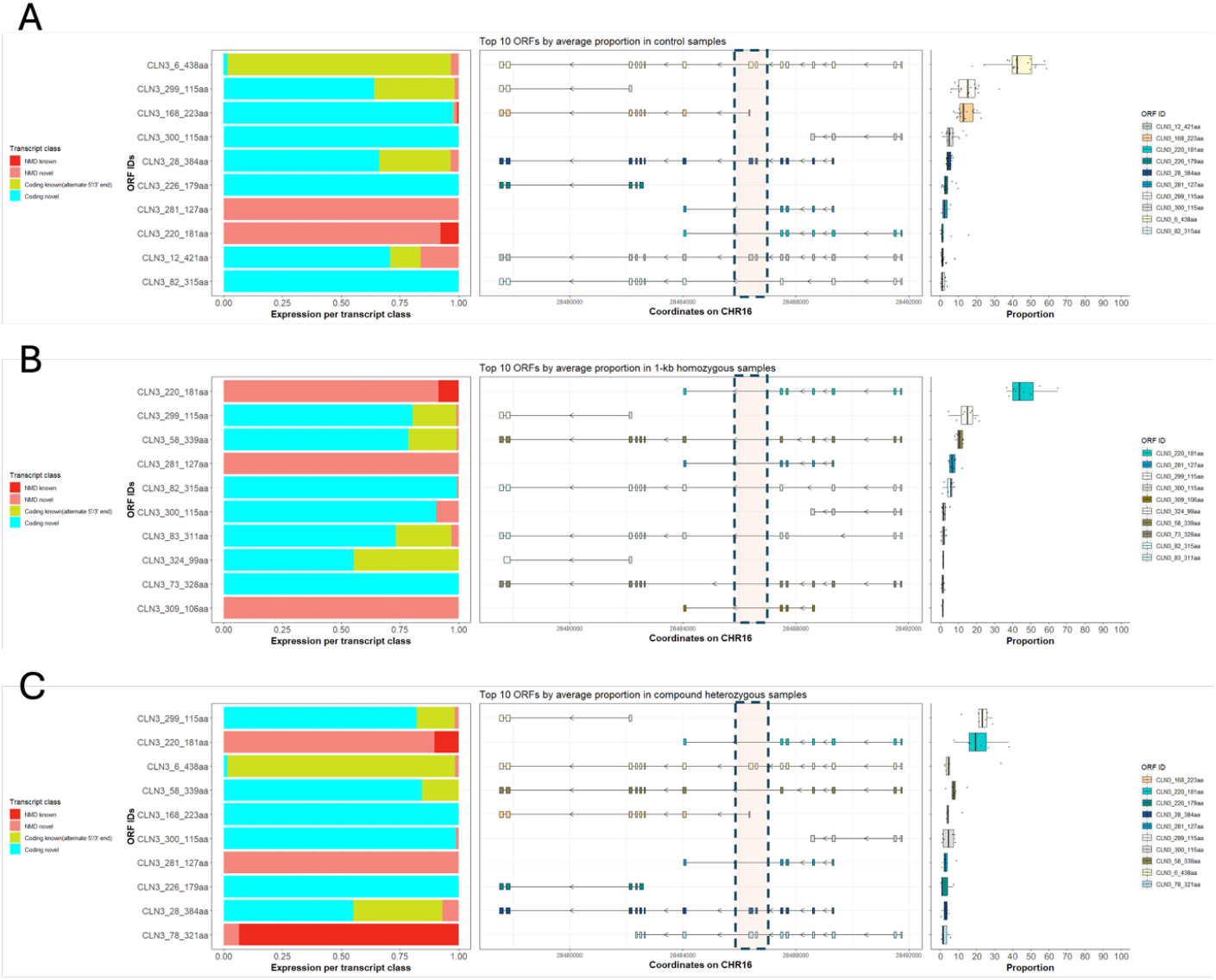
Overview of CLN3 average open reading frame expression in controls versus 1-kb deletion patient samples. A-C: From left to right, graphical representations of the top 25 transcripts in controls (A), homozygous patients. (B) and compound heterozygous patients (C), samples ranked by proportional usage in normalised expression, by transcript class (left panel), the ORF structure (middle panel) and the proportional expression of these ORF (right panel). Approximate coordinates of the 1-kb deletion are indicated by salmon boxes.

15 of the top 25 *CLN3* transcripts are coding novel, and six represent more than 2.5% each by median proportion (**Fig 2A**). These novel coding transcripts encode four of the top 10 ORF (**Fig 3A**), of which three each represent more than 10% ORF by proportion, including CLN3-223aa, observed in all individuals, accounting for a mean proportion of 13.9% (range: 22.6%-7.5%), and CLN3-384aa, with a start site in codon 8 and representing a mean proportion of 4.83% (range: 7.24-3.17%). Two sets of transcripts encode different short 115 amino acids ORFs: 3’CLN3-115aa, encoded by the 3’end of *CLN3* (exons 13-15) accounts for a mean proportion of 15% (range: 32.6-5.88%), and 5’CLN3-115aa, observed in all individuals, encoded by the 5’ end of *CLN3* (exons 1-4), accounts for a mean proportion of 6% (range: 14.2 – 0.9). Notably, several transcripts lack internal exons, including exons 7 and 8. This is discussed in detail below. The top four transcripts and top five ORF are found in all control blood samples (**Suppl Fig 3**).

### The 1-kb homozygous patients have striking and consistent effects on *CLN3* transcription, including 86 unique transcripts encoding 26 unique open reading frames

The consequence of the 1-kb intragenic deletion on transcription of *CLN3* is studied in eight patients homozygous for this deletion. This deletion, which removes exons 7 and 8 from the genome, has striking and consistent effects on *CLN3* transcription across these individuals. 86 unique transcripts are detected in homozygous patients (**Graphical figure**), of which only 5 (5.8%) transcripts are classed as coding known, 30 (34.9%) transcripts are classed as coding novel, 51 detected transcripts are predicted to undergo NMD, of which 2 (2.3%) are classed as known NMD transcripts and 49 (57%) as novel NMD transcripts.

The findings for *CLN3* transcription in 1-kb homozygous patients are summarised for the top 25 transcripts and top 10 ORFs in **Figs 2B** and **3B**, respectively. As expected, all transcripts from these homozygous patients lack exons 7 and 8 (**Fig 2B**), and so the canonical ORF CLN3-438aa is not represented. The first notable observation is the presence of multiple transcript isoforms encoding the same 181 amino acid ORF (CLN3-181aa) but differing in their 5’UTRs; they represent 10 of the top 25 transcripts (**Fig 2B**) with seven of these observed in all individuals. Together, they make up 38% of all detected transcripts (33 out of 86) and collectively result in a proportional average of 46.6% (range: 64.9-36.7%) for ORF CLN3-181aa (**Fig 3B**). The highest expressed transcript, CLN3_220_181aa_5UTR_561_3UTR_294, has a median usage of ∼13% (range: 15.6 – 7.5). These transcripts are predicted as NMD and only one is previously known. The ORF CLN3-181aa is equivalent to that encoded by the previously reported ‘major’ transcript found in patients [13].

There are several other novel transcripts occurring at high levels that are not previously reported in patients homozygous for the 1-kb deletion. Some of these encode considerably longer ORF than CLN3-181aa. Two that differ in their 5’UTRs encode the same 339 amino acid ORF (CLN3-339aa), are detected in all individuals (**Fig 2B** and **Suppl Fig 3B**), and together represent 4.7% of all detected transcripts in homozygous patients and a proportional average of 10.7% (range: 12.8 – 8.0) for ORF CLN3-339aa. Prediction algorithms suggest that these transcript isoforms are translated. The long ORF encoded by this transcript is the result of skipping exon 5 on the background of the deletion of exons 7 and 8, thereby retaining canonical sequence from exon 4, gaining non-canonical coding sequence (i.e. ORF from an alternate translation frame) for exon 6, before coming back into frame with canonical sequence in exon 9.

The previously described ‘minor ‘transcript is not in the top 25 of transcripts in 1-kb homozygous samples although its encoded long ORF CLN3-328aa is in the top 10 ORF (**Fig 3B**). This/ese together represent 1.2% of all detected transcripts and a proportional average of 1.1% (range: 2.2 - 0.2%) for ORF CLN3-328aa. The equivalent ORF has been previously modelled in yeast where it exhibited complex functional effects [15].

Other ORFs present at >5% are CLN3-315aa, CLN3-127aa and 3’CLN3-115aa and these ORF are detected in all individuals (**Suppl Fig 3B**). CLN3-315aa is a result of transcripts skipping both exons 4 and 5 causing the reading frame of exon 9 to be in frame with that of exon 3. These together represent 2.3% of all detected transcripts and a proportional average of 5.2% (range: 7.9 – 1.2%) for ORF CLN3-315aa. This ORF is also seen in controls (**Fig 2A**) but at a significantly lower expression (1.6%, range: 4.0 – 0.3). The smaller ORF CLN3-127aa is caused by transcripts that have a later start site in exon 3 and earlier stop site in exon 9 (with the novel 28 amino acids), retaining exons 4, 5, and 6. This ORF has a significant increase in mean expression in patients, being 7.1% (range: 12.2-4.6), compared to 2.8% (range: 5.6 - 0.6) in control samples. The 3’CLN3-115aa ORF maintains its relative proportional expression at a mean 14.3% (range: 21.5 - 4.6) in patients, in line with its expression in control samples.

CLN3-311aa arises due to transcripts skipping exons 3 and 4 on the background of the 1-kb deletion of exons 7 and 8. These are present at 2.3% of all detected transcripts and a proportional average of 1.9% (range: 3.5 – 0.3%) for ORF CLN3-311aa. Transcripts encoding this ORF are not present in controls but are present in 1-kb heterozygous samples, at 1.3% (range: 2.7 - 0.4).

### Previously identified disease-associated *CLN3* transcripts are present in healthy control samples

The 1-kb deletion that underlies most cases of juvenile CLN3 disease leads to many transcripts lacking exons 7 and 8. However, several transcripts lacking exons 7 and 8 are also found in all 16 of the control samples that are known not to be carriers of the 1-kb deletion (**Fig 2A**). These include seven isoforms of the so-called ‘major’ transcript (with alternative 3’/5’UTRs) encoding a 181 amino acid ORF, CLN3-181aa, of which only one was previously annotated *(ENST00000568422*.*6*) [18]. ORF CLN3-181aa is the 8^th^ highest ORF in controls with an average proportional expression of 2.57% (range: 15.7-0.25). Two isoforms of the so-called ‘minor’ transcript that additionally lacks exon 9 (with alternative 3’/5’UTRs) encode CLN3-328aa, though this ORF is not in the top 10 for control samples.

There are also other ORF encoded by transcripts lacking exons 7 and 8 in control samples. These ORF include CLN3-223aa encoded by transcripts present in all samples, two different 115 amino acid ORFs (3’CLN3-115aa and 5’CLN3-115aa) in 11 samples, CLN3-315aa in four samples, and CLN3-127aa in two samples. CLN3-315aa lacks sequence encoded by exon 4, as well as exons 7 and 8, and has an average proportional expression of 1.6% (range: 4.0-0.32). CLN3-127aa uses an alternative start site on exon 3 and is present with a mean proportion 2.84% (range: 5.62-0.62) in controls, compared to 7.08% (range: 12.2-4.59) in patients. 3’CLN3-115aa maintains its relative proportional expression at a mean 14.3% (range: 21.5-4.6) in patients in line with its expression in control samples.

Alternative splicing of variant-carrying disease genes has been recognised to reflect what is happening at a lower frequency normally [19, 20]. Here, our high-quality data allows us to quantify this for *CLN3*. Although expressed at relatively low levels in healthy controls, the highest expressed alternative ‘major’ disease transcript, CLN3_220_181aa_5UTR_561_3UTR_294, is observed 30 times in healthy samples with an average proportion of 2.2% (range: 0.33-8.49) across 5 different samples, which brings it into the top 25 transcripts (**Fig 2A**) and top 10 ORFs (**Fig 3A)** of controls. The transcript encoding ORF CLN3-328aa is observed at a lower proportional transcription of 0.72% (range 0.25-1.1) in three samples only. It is therefore not present in the top 25 transcripts or top 10 ORF in healthy controls. However, when present, it is at similar levels in patients to controls (**Fig 2B**), at 0.7% in controls and 1.1% in 1-kb homozygous patients.

We can readily compare the proportion of specific ORFs shared between patients homozygous for the 1-kb deletion and controls (**Fig 4**), and also with those that carry the 1-kb deletion *in trans* with different variants. This reveals that CLN3-181aa is expressed nearly 50-fold higher in patients than controls, CLN3-339aa is 10-fold higher, CLN3-127aa is 6-fold higher, and CLN3-223aa is 10-fold lower. For these ORFs, the proportion in heterozygous patients falls in between, as expected.

**Figure 4:**
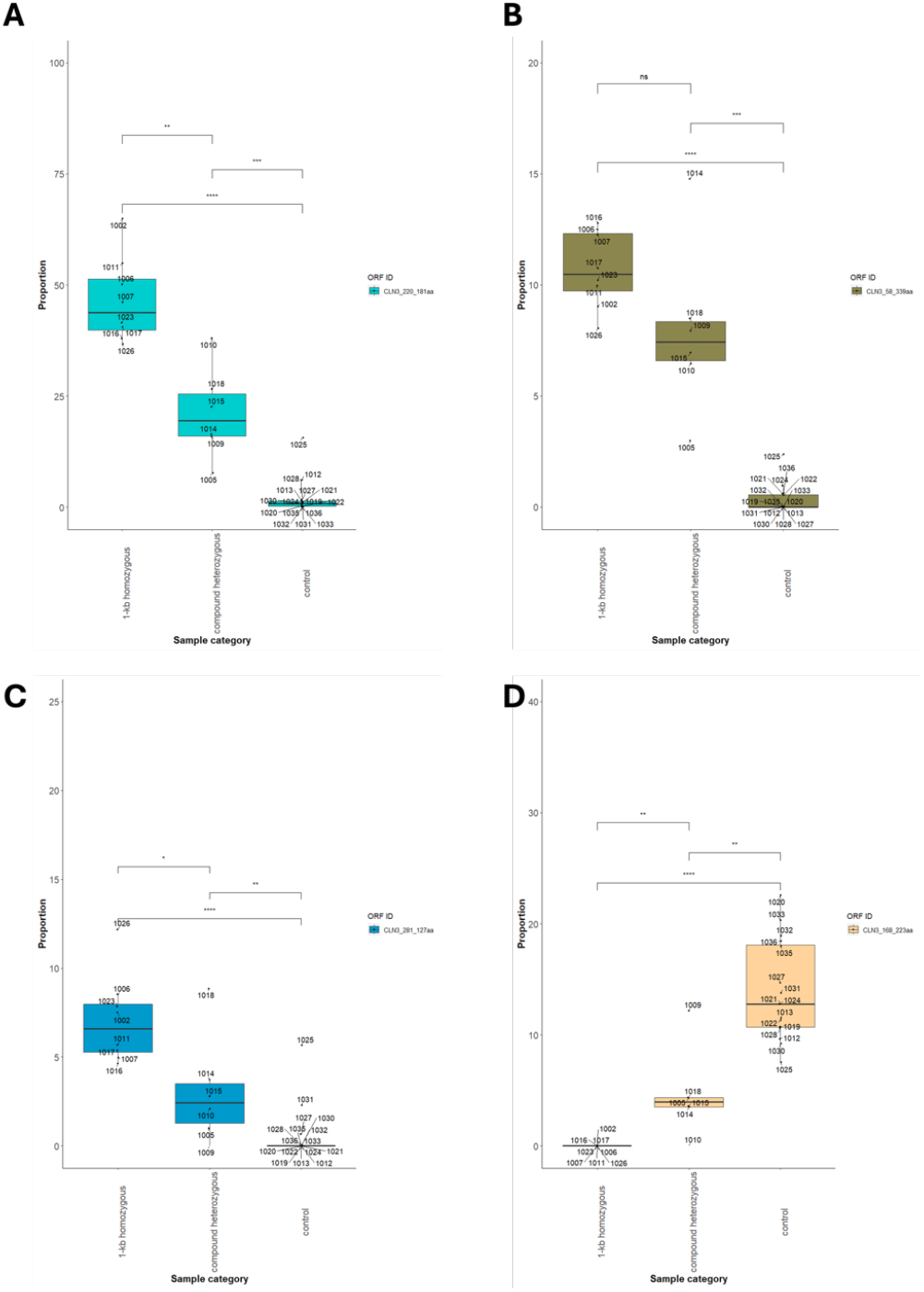
Boxplots of proportional ORF comparisons between patient and control subgroups for CLN3-181aa, CLN3-339aa, CLN3-127aa and CLN3-223aa. A-C) noticeable fold change increases across these ORFs within 1-kb homozygous patients and controls, with compound heterozygous predictably placed in between. D) Shows the decrease of CLN3-223aa within homozygous and compound heterozygous patients compared to controls.

Notably, for control individual 1025 the proportional values of ORFs more closely resembles the proportion of ORFs CLN3-181aa, CLN3-329aa, and CLN3-127aa found in heterozygotes for the 1-kb deletion. Examining the top 25 transcripts in detail from this individual (**Fig 5C**) shows many consistent with those arising from the 1-kb deletion. Together, these data suggest that this individual is a carrier for the 1-kb deletion, which is plausible given they are a member of a CLN3 disease family.

**Figure 5:**
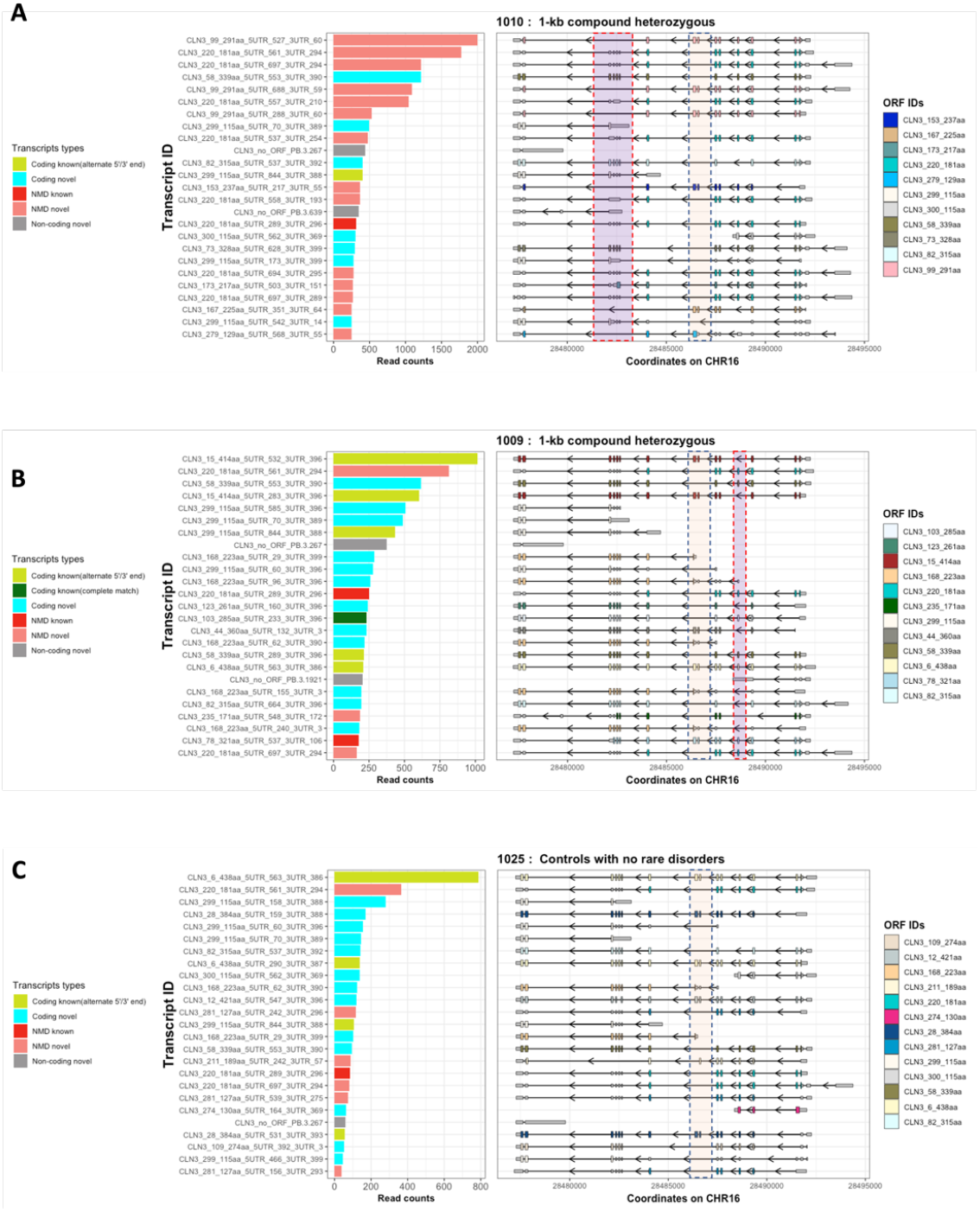
Summary of consequential changes in top 25 transcriptional structures expressed in individuals for unique mutations. A) Patient 1010 compound heterozygous for 1kb/2.8kb deletion, salmon box covers the exons Δ78 for the 1kb deletion and purple box covering the loss of Δ10-13 exons. B) Patient 1009 compound heterozygous (c.(460+1_461-1)_(677+1_678-1)del / c.290C>G, salmon box covers the 1kb-like deletion for exons Δ78 and purple boxes shows the exon skipping in exon 4. C) Control 1025 unknown mutations, but predicted from the transcriptional profile to be a 1-kb carrier, salmon box covers the exons Δ78 for the 1kb deletion. These data represent the top 25 transcripts of an individual so include non-coding transcripts.

These findings in fresh blood correlate with our previous observations of *CLN3* transcripts in public databases (www.encodeproject.org) [16, 21]. We also observed transcripts encoding CLN3-181aa in four of the commercial RNA tissue samples though not in blood (**Suppl Fig 1B’’**). Other transcripts missing exons 7 and 8 in commercial RNA tissues include those encoding 3’CLN3-115aa, CLN3-274aa and CLN3-127aa (**Suppl Fig1 B’**).

In this study, the transcripts encoding CLN3-181aa are predicted as NMD. Our experimental approach captures mature RNA with polyA tails which suggests that these transcripts may not be degraded but translated into a protein with function in both patients and healthy controls. We therefore interrogated mass spectrometry databases for unique peptide sequences of this 181 amino acid ORF [22]. We found four entries for the unique peptide sequence ‘AQPGSPS’ that is encoded by the tail end of the truncated 181 amino acid mutant protein within the Global Proteome Machine and Database (GPMDB), which is consistent with low expression in controls. Combined with its previous detection in brain tissue from multiple sclerosis (MS) patients in our earlier public data mining [16], we propose that transcripts encoding CLN3-181aa either are incorrectly predicted to undergo NMD or have properties that allow their escape from NMD. This hypothesis, if verified, is a paradigm shift in the contextualisation of juvenile CLN3 disease. We also examined the mass spectrometry databases for other CLN3 ORFs and detected sequences for CLN3-438aa and CLN3-389aa which are predominantly wildtype isoforms.

### The effect on transcription of rare disease variants are distinguishable from those of the 1-kb deletion

All eight patients homozygous for the 1-kb deletion have the same overall pattern of transcripts (**Fig 2B)**. Thus, our work demonstrates that a single individual patient provides sufficient detail and consistency for insight into transcript variation of a single gene when targeted for that gene. This is highly significant for the study of the effect of rare disease-associated variations as, for a given variant, a blood sample may only be available from one individual. We therefore explored whether it is possible to distinguish the effects on transcription of *CLN3* in individuals in this study carrying other variants to the 1-kb deletion. Six of the CLN3 disease patients in this study are carrying other variants (**Suppl Tables 2** and **3**). Two unrelated individuals carry the same variant and the 1-kb deletion, and three each carry unique variants and the 1-kb deletion. One patient carries a unique variant of a deletion removing the same exons as the 1-kb deletion.

57 unique transcripts are detected in these combined individuals (**Graphical figure**). The top 25 transcripts and top 10 ORF detected in the combined six 1-kb deletion heterozygous patients are displayed in **Figs 2C** and **3C**. The many transcriptional effects of the 1-kb intragenic deletion are readily identified due to the absence of exons 7 and 8 in transcripts and ORF. Examining only those transcripts and ORF missing exons 7 and 8, a similar pattern to those homozygous for the 1-kb deletion is observed. The highest proportional expressed transcripts and ORF are those encoding CLN3-181aa and the transcript that additionally skips exon 5, encoding CLN3-339aa. There are 13 transcripts identified overall that contain exons 7 or 8 in compound heterozygotes, although only six such transcripts in the top 25 of these grouped samples and three in the top 10 ORF. These represent the combined effects of the other disease variants. Several transcripts encoding only C terminal exons (3’CLN3-115aa) are also present in significant proportions, as in control and patient samples homozygous for the 1-kb deletion.

To determine the particular effect of a rare variant on the splicing of *CLN3* we examined data from each respective individual (**Fig 5** and **Suppl Fig 4)**. We discovered that, like the 1-kb deletion, some of the variants change the overall pattern of transcription, and this had not been predicted for this variant by programs such as AI splice. Out of these six additional disease-causing variants, only two were found not to have effects on splicing (**Suppl Table 4**).

Patient 1010 carries a large 2.8-kb deletion (c.790+352_1056+1445del) removing exons 10 to 13 in compound heterozygosity with the 1-kb deletion. The predicted effect of this deletion on translated protein is listed as p.(Gly264Valfs*29) in the NCL mutation database [23]. The effect of this deletion on transcription can be readily distinguished from that of the 1-kb deletion (**Fig 5A**), exemplified in the top two transcripts which encode the ORFs CLN3-291aa from the 2.8-kb deletion and CLN3-181aa from the 1-kb deletion. In the top 25 transcripts for this individual, six have no exons 10-13 and are derived from the 2.8-kb deletion. These encode ORFs (CLN3-291aa, CLN3-237aa, CLN3-225aa, CLN3-129aa) resulting from splicing of exon 9 to exon 14 and the introduction of a stop codon in an alternate translational frame of exon 14; their varying sizes reflect alternate start positions or splicing out of exon 4. The other 19 transcripts have no exons 7 and 8 and are like those in individuals homozygous for the 1-kb deletion.

Patient 1009 is compound heterozygous for the single nucleotide change c.290C>G in coding exon 4 which is predicted to cause a missense change, p.(Thr97Arg), and the deletion c.(460+1_461-1)_(677+1_678-1)del that removes exons 7 and 8, like the 1-kb deletion. The effect of c.290C>G and this 1-kb-like deletion on transcription can be readily distinguished (**Fig 5B**), exemplified in the top two transcripts which include one arising from the disease allele containing c.290C>G as it contains exons 7 and 8 (encoding ORF CLN3-414aa) and one from the 1-kb deletion as it lacks exons 7 and 8 (encoding ORF CLN3-181aa) (**Fig 5B**). Out of the 25 top transcripts, twelve transcripts contain exons 7 and 8 (or part thereof), so represent those deriving from the non-deletion allele. Three of these transcripts also skip exon 4 (and encode CLN3-414aa and CLN3-360aa); only transcripts encoding CLN3-414aa ORF are detected in only three of the 16 control samples. This suggests a significant effect of a change of nucleotide c.290 on splicing of the exon in which it is located. Ten transcripts are missing exons 7 and 8, giving transcripts similar to arising from the 1-kb deletion.

In contrast, patient 1005 carries variant c.512C>T in exon 7, in compound heterozygosity with the 1-kb deletion. This single nucleotide change variant is predicted to cause a simple missense change, p.(Ser171Phe). The effect of this deletion on transcription can be readily distinguished from that of the 1-kb deletion (**Suppl Fig 4**), with the top two transcripts derived from c.512C>T as they contain exons 7 and 8. In the top 25 transcripts for this individual, 11 transcripts contain exons 7 and 8 or parts thereof, and so must be derived from the c.512C>T disease allele. These encode the canonical CLN3-438aa, and ORFs CLN3-384aa, CLN3-321aa, CLN3-367aa, CLN3-223aa and 3’CLN3-115aa. These non-1-kb deletion transcripts mirror the CLN3 transcripts observed in control samples, suggesting that the effect of change of the nucleotide c.512 does not affect splicing.

Two unrelated patients (1015 and 1018) both carry c.631C>T in heterozygosity with the 1-kb deletion. This single nucleotide change variant is predicted to introduce a stop codon in coding exon 8, p.(Gln211*). The effect of this point mutation on transcription can be readily distinguished from that of the 1-kb deletion (**Suppl Fig 4**). For both individuals, several transcripts in the top 25 contain exons 7 and/or 8 and are therefore derived from the c.631C>T disease allele. These include transcripts encoding canonical CLN3-438aa (which will include the stop codon in exon 8) and CLN3-390aa (splices out only exon 7) which are observed in control samples, and a transcript encoding CLN3-373aa (splices out exon 8) that is not observed in controls. Other transcripts show structures observed in controls but not in 1-kb deletion homozygotes; these encode CLN3-223aa and CLN3-148aa. In contrast, some transcripts observed in controls were not observed in the top 25 transcripts of these two individuals, such as that encoding CLN3-384aa. Of note, each individual showed unique transcripts not readily observed in controls or individuals homozygous for the 1-kb deletion, suggesting that these may be caused by variant c.631C>T.

Patient 1014 is compound heterozygous for c.558_559del in coding exon 8, predicted as p.(Gly187fs) frameshift [23], and the 1-kb deletion. Transcripts arising from this deletion of two bases can be readily distinguished from that of the 1-kb deletion due to the presence of exons 7 and/or 8 (**Suppl Fig 4**). In the top 25 transcripts for this individual there are three transcripts containing exons 7 and 8 that encode CLN3-438aa, CLN3-384aa, CLN3-381aa, and one containing part of exon 8 that encodes CLN3-223aa. These resemble the transcript structures of controls. This suggests that deletion of these two nucleotides 558-559 in exon 8 does not affect splicing. Other transcripts in this individual not containing exons 7 and/or 8 are the same as structures observed in controls or individuals homozygous for the 1-k deletion.

Together, these analyses show that it is possible to distinguish the effects of variants from the 1-kb deletion, and also that it is possible to determine if a rare variant found in one individual affect splicing or not. The effect on transcription on *CLN3* of a deletion that removes one or more exons can readily be distinguished from wild-type transcription and the effects on transcription of the 1-kb deletion, as can that of a single nucleotide variant that affects splicing, even if this is not anticipated from the DNA change. A variant that does not affect the splicing pattern results in a set of transcripts that resemble those observed in control individuals. This is because our approach examines RNA structure, not sequence. The availability of this new type of information can be highlighted in the NCL mutation database [24] as data emerges from analysis of long-read RNA sequencing of new disease variants.

## Discussion

This study applied targeted long read RNA sequencing on blood as a representative tissue for transcript analysis of *CLN3*. We identified 679 valid *CLN3* transcripts in our blood samples, with the most highly expressed transcript accounting for only ∼28% of the total *CLN3* expression in controls. 253 of these transcripts are predicted to encode protein with novel peptide sequences due to alternate splicing. This is a new perspective on *CLN3*, and impacts our understanding of CLN3 disease. Recent research has mainly focused on describing the molecular changes and altered pathways in various cell and animal models, for example, leading to recently identified potential biomarkers for CLN3 disease [25-28]. There may still be much to uncover about the functional location(s) and mechanism of action of CLN3.

The many predicted novel coding variant transcripts from the wild-type *CLN3* locus that have not previously been annotated to human genome assembly GRChg38 suggests that these undergo translatability into different functional isoforms. Some transcripts are predicted to undergo NMD but as these are highly expressed, they may also result in novel CLN3 protein isoforms. Further exploratory research on these isoforms is required to understand their relevance and to confirm potential for translation into stable protein isoforms. Transcripts encoding the small ORF 3’CLN3-115aa are particularly interesting as these show a highly consistent expression pattern in both controls and patients (and across the different commercial tissues). This transcript, or the translated small ORF, may be involved in regulatory processes.

Taking advantage of most patients with CLN3 disease being homozygous for the 1-kb intragenic deletion, this study was able for the first time to examine the effects of this deletion on transcription at the *CLN3* locus. The findings are consistent across eight individuals and there are several striking observations. One is the increased expression of transcripts predicted to be NMD, especially the previously described ‘major’ transcript lacking exons 7 and 8 that encodes ORF CLN3-181aa. The presence of ORF CLN3-181aa is increased by ∼50-fold in these patients compared to control individuals. This high level of expression combined with the mass spectrometry detection of peptide fragments unique to the encoded CLN3-181aa in public mass spectroscopy databases are consistent with this transcript isoform not undergoing NMD but being translated into protein. The same may be true for other diseases where variant transcripts are predicted to undergo NMD, and especially those that are highly expressed. This suggests that current NMD prediction tools may not be accurate.

The observation of isoforms in patients predicted to encode longer alternative CLN3 isoforms than CLN3-181aa, including CLN3-339aa, CLN3-328aa, CLN3-315aa, and CLN3-311aa, is also striking.

These longer disease isoforms are intriguing as they may retain more substantial functionality than CLN3-181aa. These findings are consistent with our early work [13] where we showed that cells from a patient homozygous for the 1-kb deletion retain some CLN3 function. We later modelled expression of CLN3-328aa encoded by the ‘minor’ transcript, the longest ORF encoded by the two transcripts known at that time, in the fission yeast *S. pombe*, demonstrating both partial loss and new functional gains for this protein isoform [15]. Further, increased expression of the mouse equivalent of CLN3-339aa was found to reduce disease burden [27, 29] which could be consistent with residual functionality of this isoform.

Notably, only a few wild-type *CLN3* transcripts became absent due to the deletion of exons 7 and 8. These include those encoding canonical CLN3-438aa, as well as CLN3-384aa, CLN3-421aa, and CLN3-233aa which has an ATG start codon in exon 8. If CLN3 disease arises only from loss of function, and assuming that these alternative wild-type ORF provide complementary isoformic functionality, it may be that augmenting with only these four isoforms is sufficient to ameliorate the effects of disease.

All eight 1-kb homozygous patients have the same overall pattern of transcripts. This demonstrates that an individual patient provides sufficient detail and consistency to provide insight into transcript variation for a targeted gene. This is highly significant for the study of the effect of rare disease-associated variations when a sample for a given variant may only be available from one individual, and this may not be present in homozygous form. We were able to describe clear transcriptional effects for six further disease variants in this study, with only two found not to have any effects on transcript structure. For some, these were unexpected. For example, the single nucleotide change c.290C>G in exon 4 had been predicted to cause a missense change, p.(Thr97Arg). We found that this single nucleotide change affected upstream splicing and increased skipping of exon 4. In contrast, the single nucleotide change, c.512C>T in exon 8 did not affect splicing, and so is likely to cause disease solely by the predicted missense change, p.(Ser171Phe), highlighting the importance of this residue in the structure or function of CLN3. Thus, many more variants may affect splicing patterns than predicted by current splicing programs (e.g. SpliceAI) [30].

The analysis used in this work produces data at the level of exon-intron structure. Thus, structural changes in transcripts can be readily picked up. Further, this opens up the possibility of using the relative proportion of a single alternative transcript, or ideally subset of alternative transcripts, as a novel biomarker. This approach could be ideal for monitoring the effect on transcript complexity following oligonucleotide therapies to modulate splicing in personalised therapeutics [7]. This is exemplified by our observations in one healthy control sample (1025) which shows transcripts consistent with those arising from the 1-kb deletion. This person has an affected sibling homozygous for the 1-kb deletion (1026) and these results suggest that this individual is therefore likely to be a carrier for the 1-kb deletion. This supports the applicability of using the relative proportion of alternative transcripts as a biomarker sensitive enough to distinguish between healthy individuals, those homozygous for the 1-kb deletion, those that are carriers or compound heterozygous for this deletion, and those that carry other variants.

The high-quality data provided through long read RNA sequencing reveals the proportional expression of transcripts, such as the low expression (≤1%) of the ‘major’ transcript encoding CLN3-181aa in control individuals without the 1-kb deletion. The concept of disease variant-associated mis-splicing reflecting alternative splicing existing at lower levels in control samples is already established [19, 31]. That is, some disease-causing genetic variants reflect naturally occurring altered interactions of the gene with the transcription machinery. Previously, repetitive *Alu* sequences had been suggested as the original cause of the genomic 1-kb deletion [32]. In CLN3 disease patients this deletion is in linkage disequilibrium, which precludes this deletion independently occurring many times [12]. Stress granules have been observed in CLN3 patient fibroblasts [33, 34], which may suggest a link between neurodegeneration [33, 34] and the increase in alternative transcripts observed here. Regulated RNA localisation, mRNA stability and translational efficiency may be other important factors to be considered for full understanding of the transcript diversity observed in disease.

Our findings are relevant for the design of personalised medicine for CLN3 disease using antisense oligonucleotide (AON) approaches. We demonstrate for the first time the high expression of the alternative transcript lacking exons 5 as well as exons 7 and 8 (referred to elsewhere as Δ578) that encodes CLN3-339aa. The 339 amino acid protein isoform has been previously documented across several mammalian species including mouse, canines and non-human primates [18]. Its relevance to disease phenotype or CLN3 functionality remains to be elucidated [18]. CLN3-339aa has been proposed as a therapeutic agent in patients as the disease burden was reduced (but not ameliorated) when its expression was increased in a Cln3 mouse model by inducing exon 5 skipping in the alternative transcript encoding the Δ78-associated CLN3-181aa [27, 29]. This raises an interesting dichotomy as our results show that a transcript encoding CLN3-339aa is clearly already expressed in patients experiencing progressive deterioration. Whether increasing the proportion of this alternative transcript in patients further would provide benefit is therefore unclear. It may be that it is the reduction of expression of the alternative transcript encoding Δ78 -associated CLN3-181aa that is beneficial in mice, and that doing similar for this, and other toxic transcripts would be more effective at reducing the burden of disease in patients. It is therefore important to determine which individual transcripts are the most important and require a therapeutic focus, for example, which are beneficial versus detrimental to cell health. This knowledge may also aid in identifying and developing new biomarkers linked to disease aetiology and progression.

This study describes the complexity of *CLN3* transcription in humans and the effects on this in disease. We show that in humans > 90% of *CLN3* transcripts are previously unidentified transcripts, highlighting the underrepresentation seen at the *CLN3* locus compared to public databases. How this compares with transcription of the *CLN3* orthologue in other species has not been studied. This is relevant when considering the use of disease models. For example, the engineered Cln3 mouse models present a protracted form of the disease [35], and the recently developed pig model is also mild [36], however, morphant *cln3* Zebrafish show a more severe phenotype [37]. Transcriptional differences between these models and humans have not been considered and may explain some differences, and going forward without this knowledge may impair preclinical studies. It may be necessary to develop disease models that more accurately represent human *CLN3* transcription.

In summary, our study provides a foundational snapshot of the complex *CLN3* transcriptional landscape, particularly highlighting the impact of the 1-kb intragenic deletion on this. By examining patients and controls, we uncovered unannotated wild-type transcript structures that may inform deeper understanding of CLN3, and support targeting disease variants via AONs or gene replacement therapies. If alternative transcript isoforms are translated into stable proteins, our understanding of the pathogenic mechanisms of classic juvenile CLN3 disease will be changed, representing a paradigm shift. Taking into account this transcription knowledge could further improve disease-relevant models and drive therapeutic advancements. In the future, there may be a need for comprehensive transcriptomic mapping in treatment planning. This may be most pertinent for slow or fast progressing patients with the same genetic variations [38], or patients with novel disease-causing variants. Overall, our findings underscore the value of using technologies like long-read RNA sequencing to deepen our understanding of disease genes and support the development of personalised therapies for CLN3 disease and other Mendelian disorders.

## Methods

### Tissue selection

An ideal tissue source to provide RNA for analysis of the consequences of genetic variation in disease genes is blood as this is easily and non-invasively sampled and can be transported at ambient temperatures, enabling purification of RNA with high RNA integrity numbers (RIN). This is suitable for *CLN3* for the following reasons: (1) lysosomal storage material and lymphocytic vacuolization is a classic early marker phenotype of CLN3 disease patients [17], so blood lymphocytes are affected by disease, (2) our previous analysis of *CLN3* transcription in tissues in public databases suggests representative transcript variation in blood and strong correlation of blood transcript variety with that of brain tissue [16], and that the largest diversity of transcripts may be seen in blood samples.

### Commercial RNA samples

PolyA or total RNA derived from six tissues were purchased from Takara Bio. Each sample contains either total RNA or polyA RNA derived from multiple healthy adult individuals of both sexes and a range of ages (**Suppl Table 1**).

### Blood collection

Blood samples were collected at various sites using PAXgene® RNA tubes (Cat No. 762165), and following the recommended protocol. These were mixed well and kept for no more that 72h at ambient temperature (approx. 21°C), then stored long term at -70°C.

### Samples

In this study we identify the transcript profile of *CLN3* in 30 blood samples (**Suppl Table 2**). These samples were collected from 14 CLN3 disease patients of which 13 patients carry the 1-kb deletion (eight are homozygous, five are compound heterozygous), and one does not (*see also* **Suppl Table 3**). We compare with 16 control samples; these consist of three healthy individuals (one we discovered is a healthy carrier of the 1-kb deletion), four patients with other NCL genetic types (*CLN1, CLN6* and *CLN13*), and eight patients with retinitis pigmentosa arising from different causative genes (*TULP1, CRB1, COQ5, FASTKD3, PRPF31*).

### RNA extraction

RNA was extracted from blood lymphocytes using the PreAnalytiX® PAXgene® Blood RNA Kit (IVD) (Cat. no. 762174). RNA concentration and RNA integrity number (RIN) was calculated using the Fentopulseâ (Agilent). All samples had an RNA Integrity Number (RIN) >7 (average was 7.7, n=30).

### Barcoding and first strand synthesis

Samples were barcoded at first strand synthesis using a unique barcode sequence developed by IDT (**IDT probes data set**). These RNA samples were then used to generate the equivalent cDNA library barcoded for unique identification per individual source enriched for mature RNA.

cDNAs specific to *CLN3* transcripts were enriched, capturing all possible *CLN3* transcript structures expressed in human lymphocytes from both control and patient samples.

First strand synthesis and reverse transcription was conducted using Takarabio SMARTer PCR cDNA synthesis kit (Cat. No. 634926), following the manufacturer’s protocol. 245 ng of total RNA was used to generate the first strand. For each sample reaction, 3.5 ml of total RNA plus 1 ml of unique barcode primers designed from IDT was combined. Following first strand synthesis and preparation of the master mix, samples were incubated for 1.5 h at 42°C and terminated at 70°C. 40 ml of TE buffer was added to dilute the newly generated cDNA strand synthesis.

### SMARTer cDNA amplification by long distance polymerase chain reaction (PCR)

cDNA was amplified using Takara Bio Advantage 2 PCR Kit (Cat. No. 639207), following the manufacturer’s protocol. The exponential phase of amplification was optimised by comparing the output from double-stranded (ds) cDNA samples amplified for different numbers of cycles using 1.2% agarose / SYBR safe gel electrophoresis. Optimal 18 cycles were determined to be sufficient and repeated for cDNA amplification. Following amplification, samples were purified using ProNex® Size-Selective Purification System (Cat. No. NG2001) following the manufacturer’s protocol.

### Purification with ProNex® Size-Selective Purification System

Transcriptional distribution of newly generate cDNA lengths was checked using Fentopulse, to ensure good quality transcript cDNA distribution prior to multiplexing. Samples were pooled together into Eppendorf tubes with a max 24 barcoded per sample run.

### cDNA capture using Integrated DNA Technologies (IDT) Xgen® lockdown® probes

Lockdown hybridization probes were designed within IDT online tool (https://eu.idtdna.com/site/order/designtool/index/XGENDESIGN). We selected and ordered 87 non-overlapping 120-mer biotinylated hybridization probes for CLN3 which provided 100% coverage across the CLN3 locus (GENOCODEv38 annotation).

We pooled equal mass of barcoded cDNA to provide a total of 1000 ng per capture reaction. Pooled cDNA was combined with Cot DNA (7.5 ml) up to 24 max barcoded samples. Following this 1.8x.

ProNex beads were added and gently mixed 10 times (pipetting), then incubated for 10 minutes at room temperature. After washing with 200 ml freshly prepared 80% ethanol twice, residual ethanol was removed. Next 19 ml Hybridization mix was added: 9.5 ml of 2x hybridisation buffer, 3 ml of buffer enhancer, 1 ml of xGen Asym TSO block (25 nmoles), 1 ml of polyT block (25 nmole) and 4.5ml of 1X xGen Lockdown Probe pool. The PacBio target Iso-seq protocol is described in protocols.io (dx.doi.org/10.17504/protocols.io.n92ld9wy9g5b/v1).

We performed overnight hybridization reactions with 1000 ng of pooled cDNA, using the Takara Bio SMARTer PCR oligo and a poly-T blocker instead of IDT’s xGen blockers. The captured cDNA was then amplified with 16 PCR cycles, using a 4-minute extension time with the NEBNext single-cell cDNA PCR primer. We assessed the amplified captured cDNA on the Agilent FemtoPulse before preparing the sequencing library with PacBio’s SMRTbell prep kit 3.0 and sequencing on a Sequel IIe platform. Multiple bead purification and quantification steps were incorporated throughout the process.

### Automate analysis of Iso-Seq data using Analysis of PacBio Targeted Sequencing (APTARS)

The analysis of targeted CLN3 PacBio Iso-seq data was performed using the same analysis pipeline as previously [42, 43]. Using the Snakemake pipeline APTARS (https://github.com/sid-sethi/APTARS) with further optimisation to ensure proper clustering of transcript isoforms from a locus with a deletion, reads with predicted ≥ 0.99 accuracy (HiFi reads) were demultiplexed, refined and clustered using IsoSeq3 4.0.0, then aligned to the GRCh38 reference genome using minimap2 2.17. cDNA_Cupcake (v 22.0.0; https://github.com/Magdoll/cDNA_Cupcake) was used to: (i) collapse redundant transcripts, using collapse_isoforms_by_sam.py (--dun-merge-5-shorter) and (ii) obtain read counts per sample, using get_abundance_post_collapse.py followed by demux_isoseq_with_genome.py.

Transcript isoforms detected were characterised and classified using SQANTI3 (v 4.2; https://github.com/ConesaLab/SQANTI3) in combination with GENCODE 44 comprehensive gene annotation. An isoform was classified as full splice match (FSM) if it aligned with reference genome with the same splice junctions and contained the same number of exons, incomplete splice match (ISM) if it contained fewer 5′ exons than reference genome, novel in catalogue (NIC) if it is a novel isoform containing a combination of known donor or acceptor sites, or novel not in catalogue (NNC) if it is a novel isoform with at least one novel donor or acceptor site.

### Post APTARS data processing

Following transcript characterisation from SQANTI3, we applied a set of filtering criteria to remove potential genomic contamination and rare PCR artifacts and obtain valid transcripts. We removed an isoform if: (1) the percent of genomic “A” s in the downstream 20 bp window was more than 80% (“perc_A_downstream_TTS” > 80); (2) one of the junctions was predicted to be template switching artefact (“RTS_stage” = TRUE); or (3) it was not associated with the gene of interest (here, *CLN3*) for at least 50 bps. Transcripts associated with both *CLN3* and *NPIPB7* loci, the *CLN3-NPIPB7* readthrough transcripts, were not considered *CLN3* transcripts. These transcripts were filtered out.

After transcript clustering, we used SQANTI3’s output of the ORF prediction, NMD prediction and structural categorisation to group identified isoforms into the following classes with the reference annotation (GENCODE). (1) Coding known (complete match) – predicted to translate, no NMD and full-splice & UTR match with the reference genome; (2) Coding known (alternate 3’/5’ end) - predicted to translate, no NMD and full-splice but with alternative UTRs to the reference genome; (3) Coding novel - predicted to translate, no NMD and not a full-splice match with the reference genome; (4) NMD known – predicted to translate, predicted to undergo NMD and full-splice match with the reference genome; (5) NMD novel – predicted to translate, predicted to undergo NMD and not a full-splice match with the reference genome; (6) Non-coding known – predicted non-coding and full-splice match with the reference genome; (7) Non-coding novel – predicted non-coding and not a full-splice match with the reference genome.

To help calculate the normalised full-length reads (*NFLRTi*), or usage, as a percentage the following formula was used, where *NFLRTi* represents the normalised full-length read count of transcript *T* in sample *i, FLRTi* is the full-length read count of transcript *T* in sample *i* and *M* is the total number of valid *CLN3* transcripts.

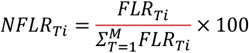

To maintain a high degree of confidence and remove potential artefacts of PCRs, we applied the following cut-offs:

1. Usage: Transcripts with usage less than 0.2% were removed from each sample.
2. Sample: When comparing the top transcripts across different sample subgroups, any transcripts not observed in at least two samples from the same subgroup were removed.

CLN3 transcripts were visualised using R Package ggtranscript [43] https://github.com/dzhang32/ggtranscript. The coding sequences used were predicted from SQANTI3.

### Prediction of Open Reading Frame (ORF)

ORFs were predicted using SQANTI3 [44], in which alternative transcripts that passed quality control and usage cut-off are collapsed down into their corresponding ORFs to observe proportional prediction of translated proteins.

### Nomenclature of transcript and ORF structures

Transcript identity follows the form ‘ORF ID-5’UTR-3’UTR’ to provide an automatic appreciation of structure; e.g. the canonical ORF has the ID CLN3_6_438aa (referred to in the general text as CLN3-438aa) and a prominent transcript has the ID ‘cln3_6_438aa_5UTR_563_3UTR_386’ meaning it has an ORF encoding a protein of 438 amino acids; all other numbers are arbitrarily assigned for ORF, 5’UTR and 3’UTR and hold consistent for this study.

This is the same pattern of nomenclature as used in our previous work [16], however only the ORF amino acid can be directly compared. The IDs for ORF, 5’UTR and 3’UTR are arbitrarily assigned in each separate study.

### Further analysis on the data

An average of 19677 full-length HiFi reads per sample (split by control n=23138 versus patient n=15152) was generated. Collapsing mapped reads resulted in the identification of 679 unique *CLN3* transcripts after quality control in blood samples.

Within the control samples, there were no unique transcripts that exactly matched annotated transcripts in GENCODE 44. We identified 132 unique *CLN3-NPIPB7* readthrough transcripts with alternative 3’/5’ untranslated regions (UTRs), as we observed previously in analysis of public long-read data [16]. These low expression level transcripts were filtered out as this work is focusing on *CLN3*.

After ranking all transcripts by relative expression for each individual sample, these were then combined by sample group. For each sample group, valid *CLN3* transcripts that passed the usage cut-off and were detected in a minimum of two samples in the corresponding group, were investigated. Collective *CLN3* transcripts detected within this study are displayed in a depreciation curve showing the number of unique transcripts (**Suppl Fig 2A, B**). There is a clear increase in the number of transcripts with low expression in CLN3 disease samples, compared to control samples with no CLN3 disease controls.

When looking at transcript profile of individuals, non-coding transcripts are included. When combining the transcript profiles of individuals by clustering their transcripts, non-coding transcript are omitted.

## Abbreviations

*NCLs*: Neuronal ceroid lipofuscinoses
*MFS*: major facilitator superfamily
*ORFs*: open-reading frames
*UTRs*: untranslated regions
*TPM*: transcripts per million
*ID*: identifier
*CDS*: coding sequence
*NMD*: nonsense-mediated decay

## Declarations

### Ethics approval and consent to participate

All UK blood samples in this study were collected with consent to participate under ethical approval via NRES Committee Bloomsbury, London (REC 13/LO/0168). Sample collection at the University Medical Center Hamburg-Eppendorf was approved by the medical ethics committees of the Ärztekammer Hamburg, Germany (PV7215). Additional international blood samples were collected under their local ethical approval in accordance with international ethical standards (Code of Ethics of the World Medical Association, Declaration of Helsinki).

### Services

UCL Long Read Sequencing Service

UCL Genomics

### Availability of data and materials

To be added

## Authors’ Contributions

Conceptualisation: C.M. and S.E.M; Methodology: C.M., H.Y.Z., E.Gn., and M.R.; Sample collection: P.M., S.E.M., C.M., G.A., N.J., A.W., A.S., M.N., N.G., F.S.; Data acquisition and analysis: C.M. and H.Y.Z, J.L., C.A..; Data interpretation: C.M., S.E.M., E.Gr, E.Gn. and M.R.; Writing – original draft: C.M. Writing – review & editing: C.M., H.Y.Z, E.Gn., J.L., C.A., E.Gr., A.S., M.N., G.A., N.J., A.W., N.G., F.S., P.M., P.G., M.R. and S.E.M.; Visualization: C.M., and H.Y.Z.; Supervision: C.M. and S.E.M.; Project administration: C.M. and S.E.M.; Funding acquisition: C.M., M.R and S.E.M. All authors read and approved the final manuscript.

## Acknowledgements

We thank families who kindly agreed to participate in research and consented to provide blood samples for research. These include, but not exclusively, families supported by the UK Batten Disease Family Association (BDFA) and NHS services, University Medical Center Hamburg-Eppendorf Germany, and a-ncl (https://www.a-ncl.it/) in Italy.

## Graphical summary figure: Overview of CLN3 transcriptional complexity and summary of transcripts identified in this study

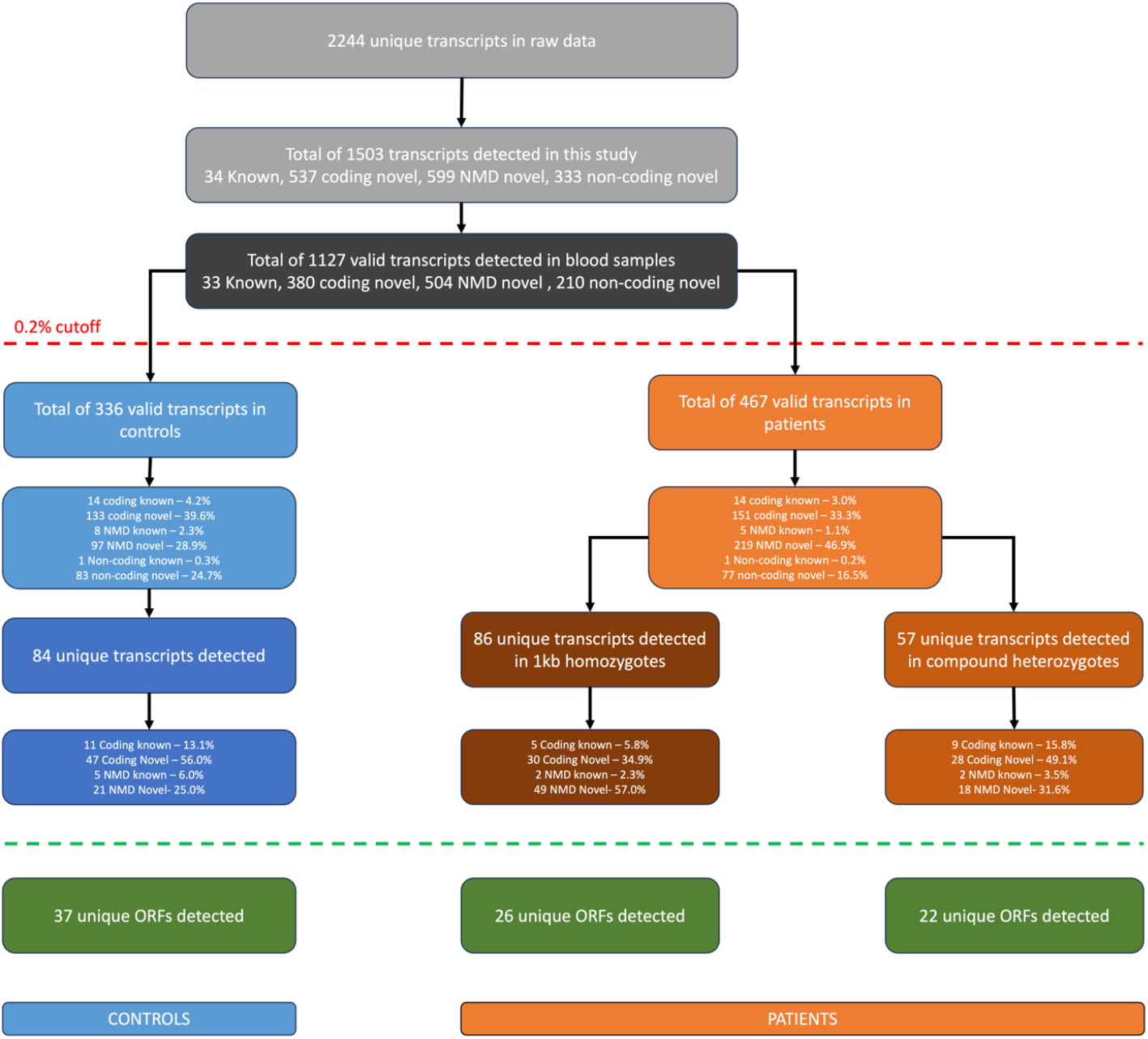

The top panel shows the number of unique transcripts present in the raw data of all samples combined and then following filtering out artefacts and read-through gene transcripts with classification (light grey boxes), and validated data extracted for the blood samples with classification (black box). To ensure confidence in the data, only those transcripts present in at least two samples and expressed at a minimum set at 0.2% normalised full-length reads (NFLR) expression across all samples were taken forward for detailed analysis (dashed red line). These middle panel summarises the analysed data for controls (light blue boxes) and patients which are subdivided into two groups (orange boxes) to show the total number of transcripts and their classifcation and including any classed as non-coding, and then the number of unique coding transcripts and their classification (dark blue and two shades of dark orange boxes). The lower panel summarises the number of unique ORFs encoded by unique transcipts in each group (green boxes). % are rounded to 1 dp so may not add up to 100%.

