## Supplementary Figures for "Targeted long-read RNA sequencing reveals the complexity of *CLN3* transcription and the consequences of the most common 1-kb deletion in patients with juvenile CLN3 disease"

### Supplementary figures /tables

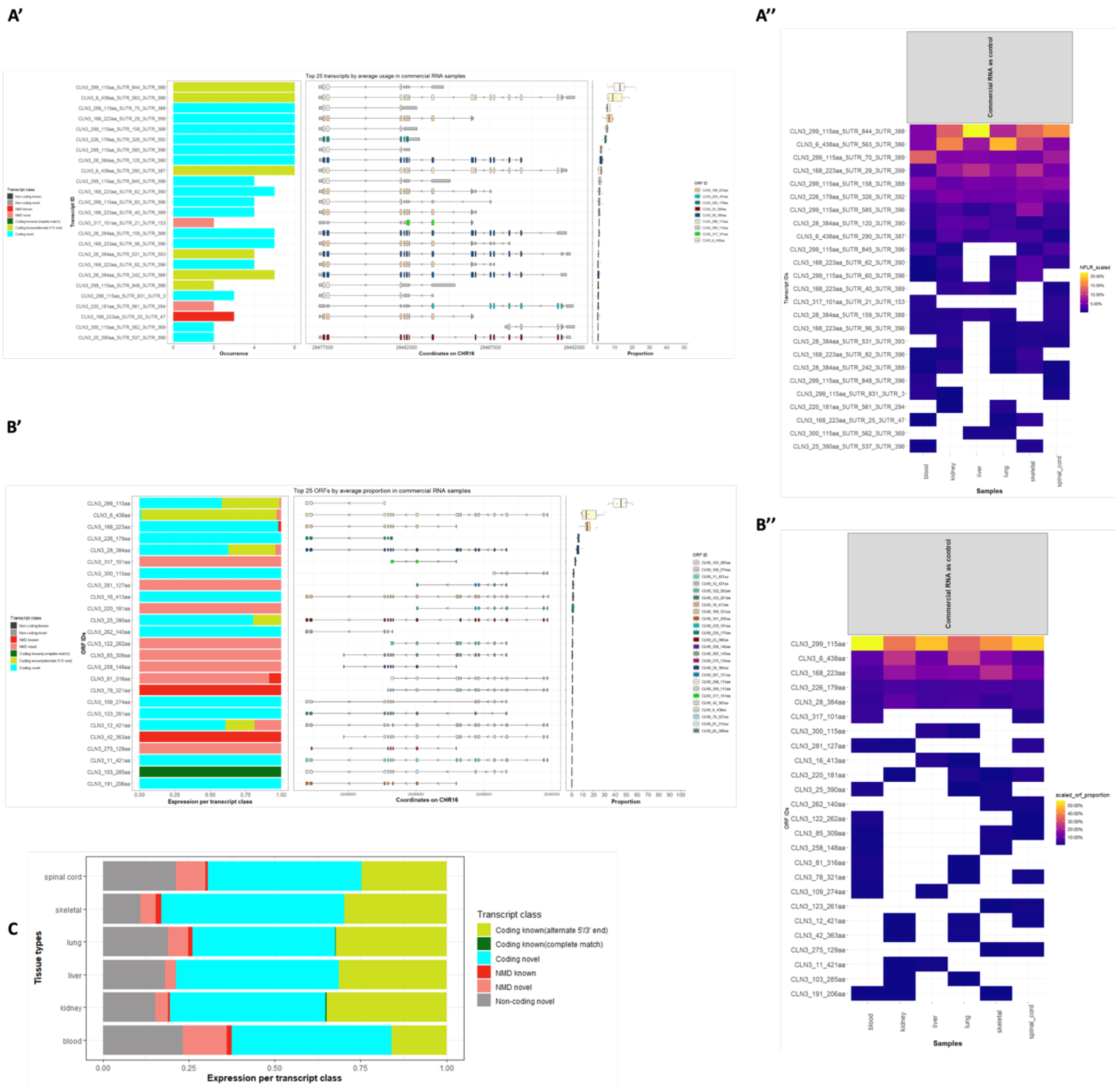


**Supplementary Figure 1**: Overview summary of *CLN3* transcript variation in commercial RNA samples from various tissues. A'-B': From top to bottom, graphical representations of the top 25 *CLN3* transcripts from all commercial RNA samples. A') by transcript occurrence (left panel), transcript structure with predicted open reading frames (ORF) (coloured extended boxed) and non-coding exons (grey, smaller boxes) (middle panel) and average transcript expression based on normalised usage (right panel). B') samples ranked by proportional usage in normalised expression, by transcript class (left panel), the ORF structure (middle panel) and the proportional expression of these ORFs (right panel). A'') heat map of the top 25 *CLN3* transcripts proportional expression levels across each specific tissues. B"") heat map of the top 25 CLN3 ORFs proportional expression levels across each specific tissues. C) Summary of *CLN3* transcript class by tissue type for commercial RNA.

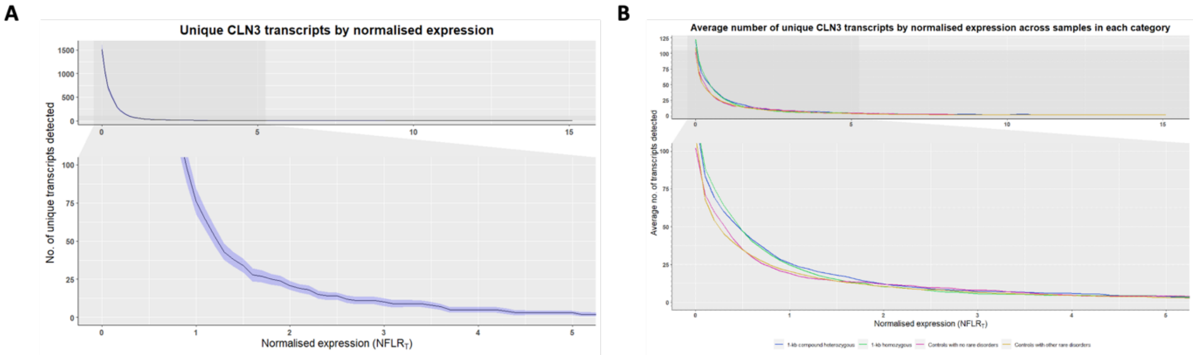


**Supplementary Figure 2**: Overview summary of *CLN3* transcriptional data.  A) Depreciation curve showing the number of unique CLN3 transcripts on the Y-axis increased by increasing the normalized full-length read count of transcript (NFLRT) on the X-axis across all samples normalised. B) The disease affects CLN3 transcription as the number of transcripts with low expression is increased in CLN3 disease samples compared to control samples with no CLN3 disease.  Average number of transcripts expressed across four subgroups, 1-kb compound heterogenous, 1-kb homozygous, healthy controls with no disease, controls with disease that is not CLN3 disease, all normalised.

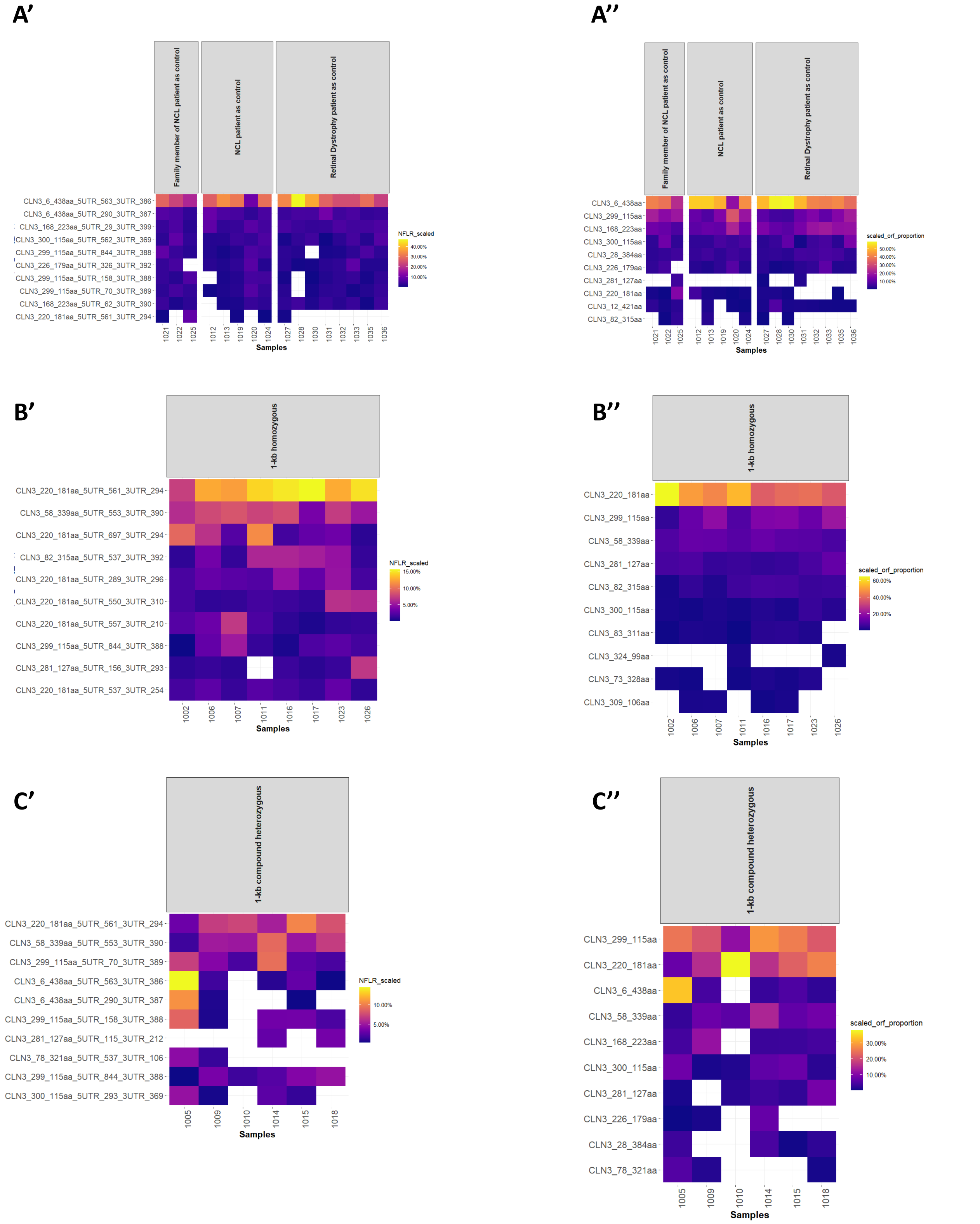


**Supplementary Figure 3**: Summary heatmap of proportional *CLN3* expression levels of the top 10 transcript structures (A'-C') and top 10 ORFs (A''-C'') for control (A), 1-kb homozygous patients (B) and 1-kb heterozygous patient (C) blood samples.

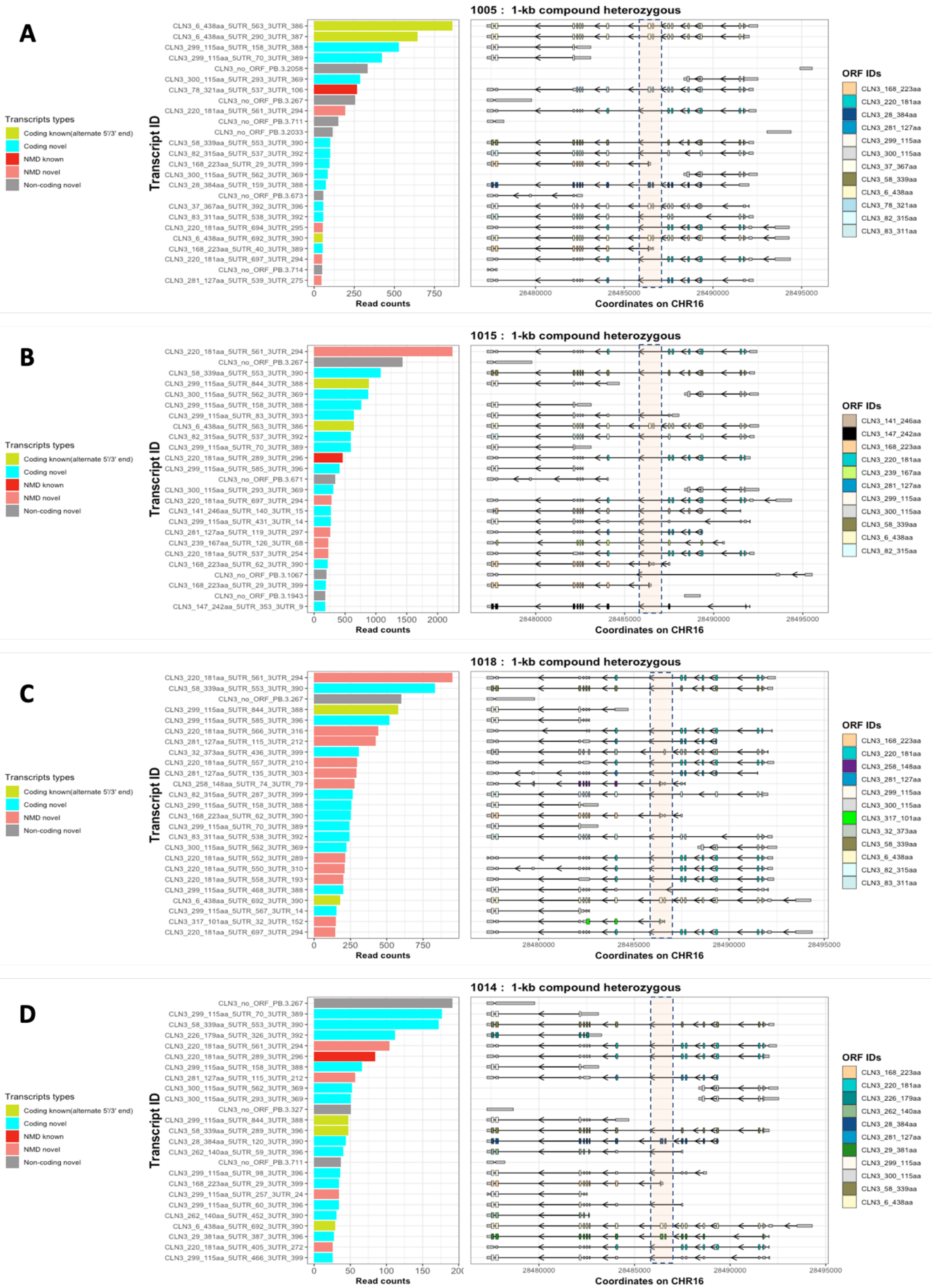


**Supplementary Fig 4**: Summary of consequential changes in top 25 transcriptional structures expressed in four individuals for their unique variants. A) Patient 1005 compound heterozygous for 1-kb deletion; c.512C>T. B) and C) Patients 1015 and 1018 compound heterozygous 1-kb deletion; c.631C>T. D) Patient 1014 compound heterozygous for the 1-kb deletion and homozygous for c.558_559del; c.558_559del. These data represent the top 25 transcripts of an individual so include non-coding transcripts.

**Supplementary Table 1: Summary of commercial RNA tissue type and how they were derived.**

| Sample barcode | Tissue Type | Type of RNA | Number and sex of pooled individuals | Age range of pooled individuals | Cause of death |
|---|---|---|---|---|---|
| 1001 | Liver | Total RNA | 3 males/ female | 64-78 | N/A |
| 1038 | Kidney | PolyA | 4 male/female | 28-48 | Sudden death |
| 1039 | Skeletal muscle | PolyA | 5 male/female | 27-60 | Sudden death |
| 1044 | Peripheral leukocytes | PolyA | Unspecified male/female | 18-40 | N/A |
| 1047 | Spinal cord | PolyA | 18 male/female | 25-63 | Sudden death |
| 1048 | Lung | PolyA | 3 male/female | 32-61 | Sudden death |

N/A is not applicable

**Supplementary Table 2: Summary of blood sample descriptions with information provided on barcode, control versus patient, characteristics and disease diagnosis, associated gene of disease and gene variations.**

| Sample barcode | Control/ Patient | Sex/Age (years) | Disease diagnosis | Disease Gene | Gene variation |
|---|---|---|---|---|---|
| 1002 | Patient | F/17 | CLN3 disease | *CLN3* | 1-kb ; 1-kb |
| 1005 | Patient | F/12 | CLN3 disease | *CLN3* | 1-kb ; c.512C>T |
| 1006 | Patient | F/17 | CLN3 disease | *CLN3* | 1-kb ; 1-kb |
| 1007 | Patient | M/15 | CLN3 disease | *CLN3* | 1-kb ; 1-kb |
| 1009 | Patient | F/8 | CLN3 disease | *CLN3* | c.(460+1_461-1)_(677+1_678-1)del ; c.290C>G |
| 1010 | Patient | F/15 | CLN3 disease | *CLN3* | 1-kb ; 2.8-kb deletion |
| 1011 | Patient | M/23 | CLN3 disease | *CLN3* | 1-kb ; 1-kb |
| 1012 | Control | M/11 | CLN1 disease | *CLN1* | c.490C>T ; c.541G>A |
| 1013 | Control | M/15 | CLN1 disease | *CLN1* | c.364A>T ; c.364A>T |
| 1014 | Patient | F/12 | CLN3 disease | *CLN3* | 1-kb, c.558_559del ; c.558_559del |
| 1015 | Patient | M/23 | CLN3 disease | *CLN3* | 1-kb ; c.631C>T |
| 1016 | Patient | F/9 | CLN3 disease | *CLN3* | 1--kb ; 1-kb |
| 1017 | Patient | F/12 | CLN3 disease | *CLN3* | 1-kb ; 1-kb |
| 1018 | Patient | M/23 | CLN3 disease | *CLN3* | 1-kb ; c.631C>T |
| 1019 | Control | F/11 | CLN6 disease | *CLN6* | c.307C>T p.(Arg103Trp) ; c.816dup p.(Val273Cysfs*11) |
| 1020 | Control | M/44 | CLN13 disease | *CLN13* | c.593_594del p.(Tyr198*) ; c.971T>C p.(Leu324Ser) |
| 1021 | Control | M/75 | Unaffected | *CLN13* | Unaffected CLN13 family member (1020) |
| 1022 | Control | F/72 | Unaffected | *CLN13* | Unaffected CLN13 family member (1020) |
| 1023 | Patient | F/11 | CLN3 disease | *CLN3* | 1-kb ; 1-kb |
| 1024 | Control | M/14 | CLN1 disease | *CLN1* | c.451C>T (p.Arg151*) ; c.739T>C p.(Tyr247His) |
| 1025 | Control | F/17 | Unaffected | *CLN3* | Unaffected CLN3 family member (1026) |
| 1026 | Patient | M/19 | CLN3 disease | *CLN3* | 1-kb ; 1-kb |
| 1027 | Control | M/66 | Retinal Pigmentosa | *TULP1* | *c.1496-6C>T ; c.1496-6C>T (published PMID 36084042)* |
| 1028 | Control | M/53 | Retinal Dystrophy | *CRB1* | c.2290C > T ; c.3879-1203C>G *(published PMID 36084042)* |
| 1030 | Control | M/55 | Retinal Pigmentosa | *COQ5* | c.933delC ; c.682-7T>G *(published PMID 36266294)* |
| 1031 | Control | F/18 | Optic Atrophy | *FASTKD2* | c.645G>C ; chr2:206764971C>T (upstream of *FASTKD2* in *MDH1B*) |
| 1032 | Control | M/59 | Unaffected | *FASTKD2* | c.645G>C |
| 1033 | Control | F/50 | Unaffected | *FASTKD2* | chr2:206764971C>T (upstream of *FASTKD2* in *MDH1B*) |
| 1035 | Control | F/63 | Retinal Pigmentosa | *PRPF31* | c.946-380T>C (dominant) |
| 1036 | Control | F/36 | Retinal Pigmentosa | *PRPF31* | c.1073+5G>A (dominant) |

F= Female sex; M = Male sex. Cells are colour-coded: Patient (yellow) vs control (grey); genetic mutation when shared with another sample – homozygous 1-kb deletion ('1-kb') is dark green with unaffected family member light green; other CLN3 variants are shades of pink; CLN1, CLN6 and CLN13 diseases including unaffected family members are shades of blue; non-NCL disease have no colour.

#### Supplementary Table 3: Summary of mutations across CLN3 patients.

| Gene variant | Barcode |
|---|---|
| 1-kb ; 1-kb | 1002, 1006, 1007, 1011, 1016, 1017, 1023, 1026 |
| 1-kb ; c.512C>T | 1005 |
| c.(460+1_461-1)_(677+1_678-1)del ; c.290C>G (p.(Thr97Arg)) | 1009 |
| 1-kb ; 2.8-kb deletion | 1010 |
| 1-kb, c.558_559del ; c.558_559del | 1014 |
| 1-kb ; c.631C>T | 1015, 1018 |

Cells are colour-coded for genetic mutation – homozygous 1-kb deletion is dark green; other CLN3 variants are shades of pink, with darker pink representing a variant deletion on one disease allele that is similar to the 1-kb deletion (predicted as p.(Ala154Thrfs*28)).

**Supplementary Table 4:** Summary of the consequences of non-1-kb deletion variants across compound heterozygous patients as predicted by Splice AI and observed in this study.

| Sample barcode | Gene | DNA | Location by coding exon | Predicted effect on protein | Splice AI prediction | Observed effects on transcript structure compared to controls |
|---|---|---|---|---|---|---|
| 1005 | *CLN3* | c.512C>T | Exon 7 | p.(Ser171Phe) | Missense variant | Like control |
| 1009 | *CLN3* | c.290C>G | Exon 4 | p.(Thr97Arg) | Missense variant | Increased splicing out of exon 4 |
| 1014 | *CLN3* | c.558_559del | Exon 8 | p.(Gly187fs) | Frameshift variant | Like control |
| 1015, 1018 | *CLN3* | c.631C>T | Exon 8 | p.(Gln211*) | Nonsense, Stop codon gained | Increased splicing out of exon 8, and other variably spliced transcripts not found in controls |
| 1010 | *CLN3* | 2.8-kb deletion | Exons 10, 11, 12, 13 deleted | p.(Gly264Valfs *29) | Splice acceptor variant | Transcripts do not contain exons 10-13 and a stop codon is introduced in exon 14 |

Exon numbering is for coding exons as for the that encoded by the canonical CLN3-438aa ORF.
Cells for barcode are colour-coded for genetic mutation as for Suppl Table 3 – CLN3 variants are shades of pink.
